# GSDMD Deficiency Protects Against Aortic Rupture

**DOI:** 10.1101/2021.01.08.425983

**Authors:** Dien Ye, Deborah A. Howatt, Zhenyu Li, Alan Daugherty, Hong S. Lu, Congqing Wu

**Affiliations:** Saha Cardiovascular Research Center, University of Kentucky, Lexington, KY; Department of Internal Medicine, University of Kentucky, Lexington, KY; Department of Physiology, University of Kentucky, Lexington, KY

## Abstract

**Objective:** Aortic ruptures are fatal consequences of aortic aneurysms with macrophage accumulation being a hallmark at the site of ruptures. Pyroptosis is critical in macrophage-mediated inflammation. This study determined effects of pyroptosis on aortic dilation and rupture using GSDMD deficient mice.

**Approach and Results:** In an initial study, male *Gsdmd*^*+/+*^ and *Gsdmd*^*-/-*^ mice in C57BL/6J background (8 – 10 weeks old) were infected with adeno-associated viral vectors encoding mouse PCSK9D377Y gain-of-function mutation and fed a Western diet to induce hypercholesterolemia. After two weeks of AAV infection, angiotensin II (AngII, 1 µg/kg/min) was infused. During the 4 weeks of AngII infusion, 5 of 13 *Gsdmd*^*+/+*^ mice died of aortic rupture, whereas no aortic rupture occurred in *Gsdmd*^*-/-*^ mice. In surviving mice, no differences in either ascending or abdominal aortic dilation were observed between *Gsdmd*^*+/+*^ and *Gsdmd*^*-/-*^ mice. To determine whether protection of GSDMD deficiency against aortic rupture is specific to AngII infusion, we subsequently examined aortic pathologies in mice administered beta-aminopropionitrile (BAPN). BAPN (0.5% wt/vol) was administered in drinking water to male *Gsdmd*^*+/+*^ and *Gsdmd*^*-/-*^ mice (4 weeks old) for 4 weeks. Six of 13 *Gsdmd*^*+/+*^ mice died of aortic rupture, whereas no aortic rupture occurred in *Gsdmd*^*-/-*^ mice. In mice survived, no differences of diameters in the ascending, arch, or abdominal aortic regions were observed between *Gsdmd*^*+/+*^ and *Gsdmd*^*-/-*^ mice.

**Conclusions:** GSDMD deficiency protects against AngII or BAPN-induced aortic ruptures in mice.

**Highlights:**

1. GSDMD deficiency protects against angiotensin II-induced aortic rupture in hypercholesterolemic mice.
2. GSDMD deficiency protects against beta-aminopropionitrile (BAPN)-induced aortic dissection and rupture in C57BL/6J mice.

## INTRODUCTION

Aortic aneurysms have high risk for aortic dissection and rupture that lead to high mortality.^1^ Most studies report aortic diameters to evaluate the disease severity. However, rupture of smaller aortic aneurysms also occurs in humans,^2-4^ suggesting aneurysm size is a limited predictor of aortic rupture.

Inflammation is profound in dissected and ruptured abdominal aortic aneurysms (AAAs).^1, 5^ Inflammatory lesions often contain dead cells. Pyroptosis, distinct from apoptosis, is a lytic form of programmed cell death.^6, 7^ This process is dependent on the cleavage of gasdermin D (GSDMD) by inflammatory caspase-1 or −11.^8-11^ Pyroptosis and production of interleukin (IL)-1*β/*IL-18 are the prominent outcomes of inflammasome activation in response to pathogenic molecules as well as danger signals and tissue damage.^12, 13^ Inflammasomes contribute to aortic aneurysms.^14, 15^ However, aortic aneurysm studies of inflammasomes focused on IL-1β have yielded conflicting results.^16-20^ Of note, the clinical trial of AAAs on an IL-1β antibody (NCT02007252) was terminated prematurely due to lack of efficacy and futility. Whether GSDMD-mediated pyroptosis contributes to aortic aneurysms, dissection, and rupture remains unknown.

In this study, we determined effects of genetic GSDMD deficiency on aortic aneurysms, dissection, and rupture in two mouse models. One was angiotensin II (AngII) infusion to hypercholesterolemic mice; the other was administration of beta-aminopropionitrile (BAPN) in young normolipidemic mice.

## RESULTS AND DISCUSSION

### GSDMD deficiency prevents AngII-induced aortic rupture

To induce hypercholesterolemia, which augments AngII-induced aortic aneurysms,^21^ we injected AAVs encoding mouse PCSK9D377Y to male *Gsdmd*^*+/+*^ or *Gsdmd*^*-/-*^ mice and fed them a Western diet for 2 weeks prior to AngII infusion.^22^ This study only used male mice because female mice have low incidence of AngII-induced aortic aneurysms.^23^ Plasma total cholesterol concentrations were increased greatly within 2 weeks in both male *Gsdmd*^*+/+*^ and *Gsdmd*^*-/-*^ mice (Figure 1A). During AngII infusion, 5 of 13 mice died due to aortic rupture in *Gsdmd*^*+/+*^ mice, but no death occurred in *Gsdmd*^*-/-*^ mice (Figure 1B). After 4 weeks of AngII infusion, survived mice were terminated, aortas were dissected and cleaned to measure surface area of the ascending and arch region (representing ascending and arch dilation) and outer width of the abdominal region (representing abdominal aortic dilation) since AngII induces dilations in both ascending and abdominal aortic regions (Figure I in the online-only Data Supplement). We did not detect differences on surface area of the ascending aorta and maximal outer diameters of suprarenal aorta between *Gsdmd*^*+/+*^ and *Gsdmd*^*-/-*^ mice (Figure 1C and D). These findings suggest that depletion of GSDMD prevents aortic ruptures, but does not affect aortic dilation.

**Figure 1.**
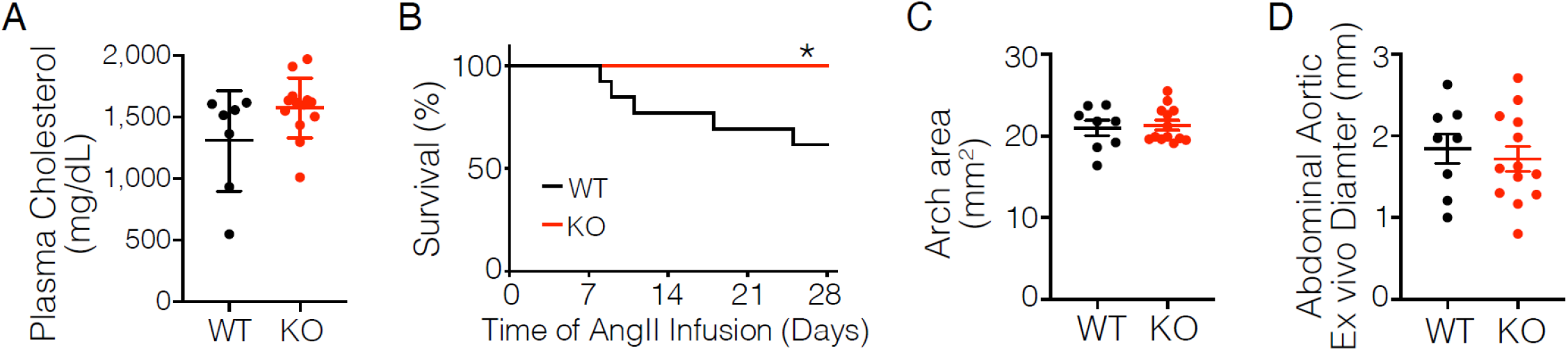
Deficiency of GSDMD prevents aortic rupture in AngII-infused mice. Male *Gsdmd*^*+/+*^ (WT) and *Gsdmd*^*-/-*^ (KO) mice in C57BL/6J background were injected intraperitoneally with AAVs encoding mouse PCSK9D377Y, fed a Western diet, and infused with AngII. (**A**). Plasma total cholesterol concentrations were measured by an enzymatic assay kit. (**B**). Survival rate of mice were analyzed by LogRank analysis. * P < 0.05. (**C**). Surface area was measured using an *en face* method. (**D**). Maximal outer width was measured using an *ex vivo* method.

### Deficiency of GSDMD prevents BAPN-induced aortic rupture

Lysyl oxidase (LOX) and LOX-like (LOXL) contribute to cross-linking of collagen and elastin of the aorta. BAPN inhibits LOX and LOXL enzymatic activity.^24, 25^ In young mice at the age of 3 – 4 weeks, BAPN induces aortic dissection and rupture.^26^ We administered BAPN to male *Gsdmd*^*+/+*^ or *Gsdmd*^*-/-*^ mice when they were 4 weeks old. Six of 13 *Gsdmd*^*+/+*^ mice died of descending aortic rupture as demonstrated by necropsy (Figure 2A), but no *Gsdmd*^*-/-*^ mice died of aortic rupture. In mice survived after 4 weeks of BAPN administration, aortic pathologies were found in the ascending, arch, and descending thoracic regions (Figure II in the online-only Data Supplement). Ascending, arch, and descending aortic dilations were not different between *Gsdmd*^*+/+*^ and *Gsdmd*^*-/-*^ mice (Figure 2B). No apparent dilation was noted in abdominal aortas of both *Gsdmd*^*+/+*^ and *Gsdmd*^*-/-*^ mice (Figure 2C). These findings are consistent with the AngII-infused mouse model that GSDMD deficiency prevents aortic rupture but not aortic dilation.

**Figure 2.**
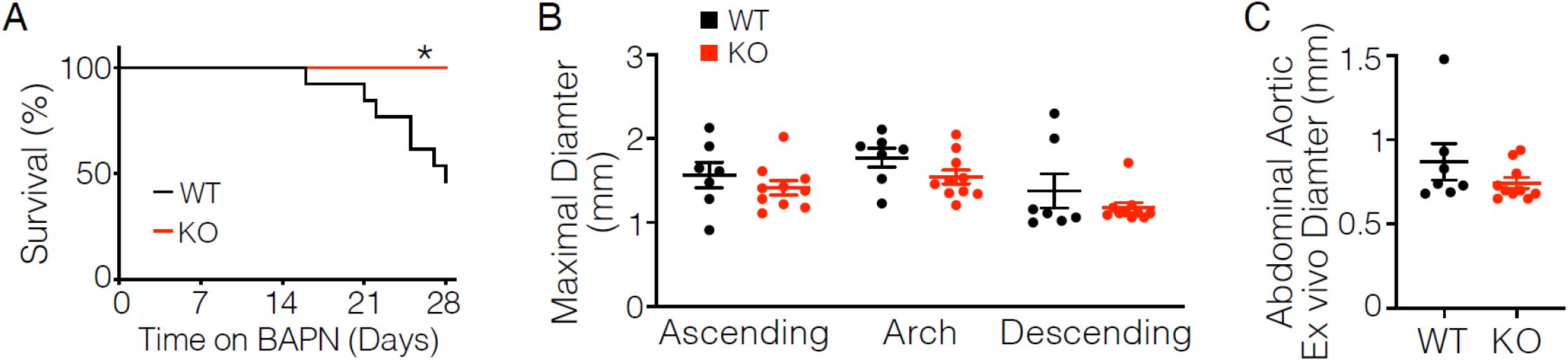
Deficiency of GSDMD prevents aortic rupture in mice administered BAPN. Male *Gsdmd*^*+/+*^ and *Gsdmd*^*-/-*^ mice in C57BL/6J background were studied. (**A**). Survival rate of mice were analyzed by LogRank analysis. * P < 0.05. (**B**). Maximal diameter was measured on in situ images. (**D**). Maximal outer width was measured using an ex vivo method.

In summary, deficiency of GSDMD, the executioner of macrophage pyroptosis, protects against aortic ruptures in two aortic aneurysm animal models: mice infused with AngII or administered BAPN. Macrophage accumulation, accompanied by elastin fragmentation, is a hallmark of aortic aneurysmal tissues.^27, 28^ Pyroptotic macrophages release cellular content in an uncontrolled manner, which could promote MMPs release into extracellular matrix, contributing to elastin fragmentation. Our data provide solid evidence that inhibition of macrophage pyroptosis by GSDMD deficiency is effective in aortic rupture prevention. Further studies are warranted to determine the molecular and cellular mechanisms by which depletion of GSDMD prevents aortic ruptures.

## MATERIALS AND METHODS

### Animals

*Gsdmd*^*-/-*^ mouse strain was provided by Dr. Toshihiko Shiroishi at the National Institute of Genetics, Shizuoka, Japan. *Gsdmd*^*+/+*^ and *Gsdmd*^*-/-*^ mice on a C57BL/6J background were bred in-house.^29^ All animal experiments reported in this manuscript were performed with the approval of the University of Kentucky Institutional Animal Care and Use Committee.

### Adeno-associated viral (AAV) vectors

AAV vectors (serotype 8) driven by a hepatocyte-specific thyroxine-binding globulin (TBG) promoter were produced by the Vector Core in the Gene Therapy Program at the University of Pennsylvania. These AAV vectors contained inserts expressing mouse PCSK9D377Y mutation (equivalent to human PCSK9D374Y gain-of-function mutation).^22^ For AngII infusion study, male *Gsdmd*^*+/+*^ and *Gsdmd*^*-/-*^ mice at the age of 8 – 10 weeks were injected intraperitoneally with AAV vectors (2 × 10^11^ genome copies/mouse), and fed a Western diet (TD.88137, Envigo) for 2 weeks to induce hypercholesterolemia.

### Osmotic mini pump implantation and angiotensin II (AngII) infusion

AngII (1 µg/kg/min; Cat# H-1706; Bachem) was infused subcutaneously via mini osmotic pumps (Alzet Model # 2004; Durect Corp.) for 4 weeks, preceded by 2 weeks of Western diet feeding. Western diet continued during AngII infusion. Surgical procedure is described in our previous publication.^30^

### Administration of beta-aminopropionitrile (BAPN)

Male *Gsdmd*^*+/+*^ and *Gsdmd*^*-/-*^ mice at the age of 4 weeks were administered BAPN (0.5% wt/vol; Cat # A3134; Sigma) in drinking water for 4 weeks. Fresh drinking water containing BAPN was changed twice a week.

### Plasma cholesterol measurements

Mouse blood were collected in the presence of EDTA (final concentration: 1.8 mg/ml) by cardiac bleeding via right ventricle. Plasma total cholesterol concentrations were measured using an enzymatic commercial kit (Cat # 999-02601; Wako Chemicals USA).

### Statistical analysis

Data are represented as means ± standard error of means (SEM). SigmaPlot version 14.0 (SYSTAT Software Inc.) was used for statistical analyses. Survival Log-Rank analysis was used to compare the cumulative survival rate. Incidence of aortic rupture was analyzed by Fisher exact test. Aortic diameters were analyzed by Student’s t-test if data passed normality and equal variance tests. If either normality or equal variance test failed, data were analyzed by Mann-Whitney rank sum test. P < 0.05 was considered statistically significant.

## ACKNOWLEDGMENTS

We thank Dr. Toshihiko Shiroishi at the National Institute of Genetics, Shizuoka, Japan for providing *Gsdmd*^*-/-*^ mouse.

## Non-standard abbreviations

AAV: adeno-associated viral vectors
AngII: angiotensin II
AAAs: abdominal aortic aneurysms
BAPN: beta-aminopropionitrile
GSDMD: gasdermin D

---

**Figure I.**
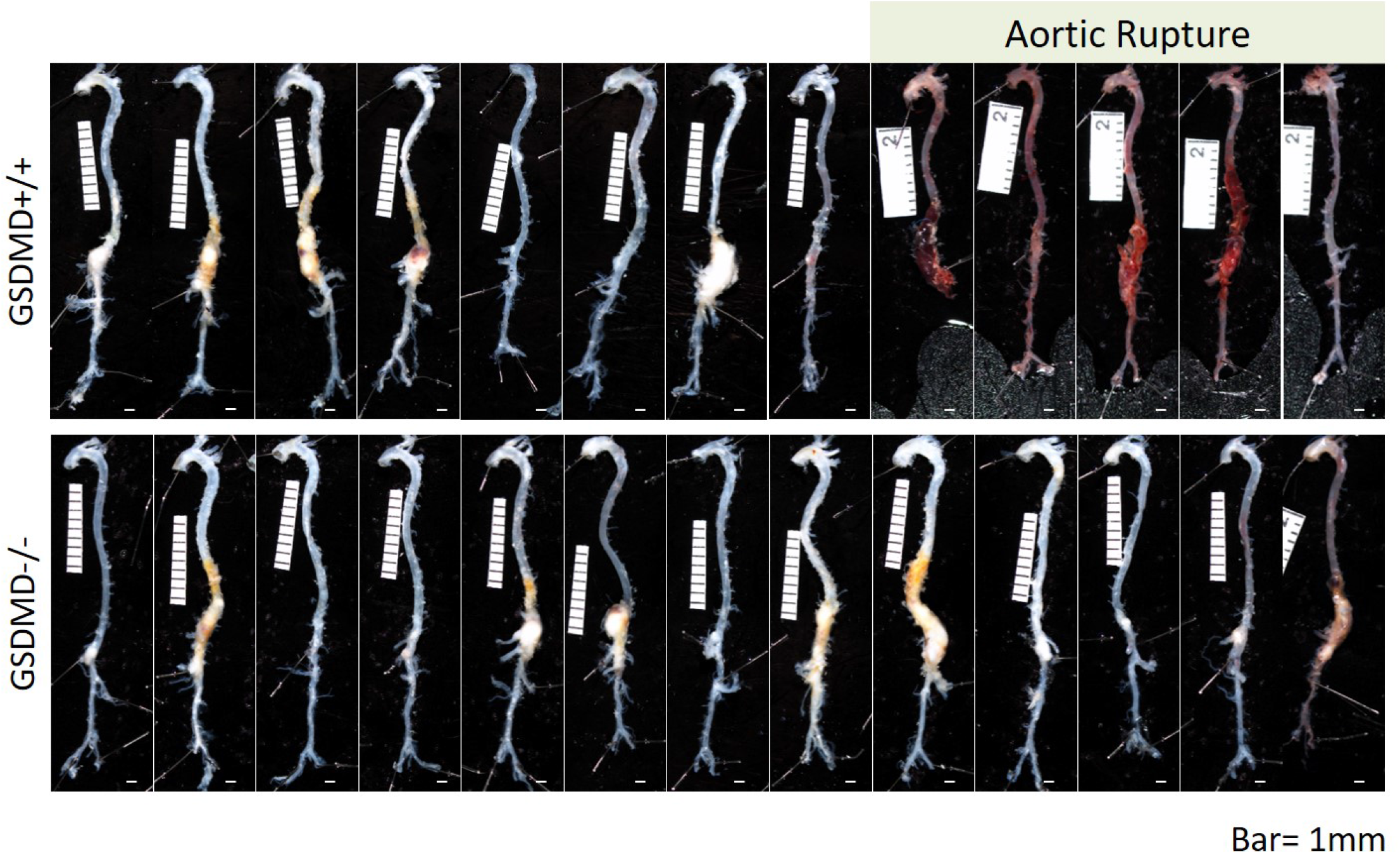
Ex vivo images of aortas from *Gsdmd*^*+/+*^ and *Gsdmd*^*-/-*^ mice infused with AngII.

**Figure II.**
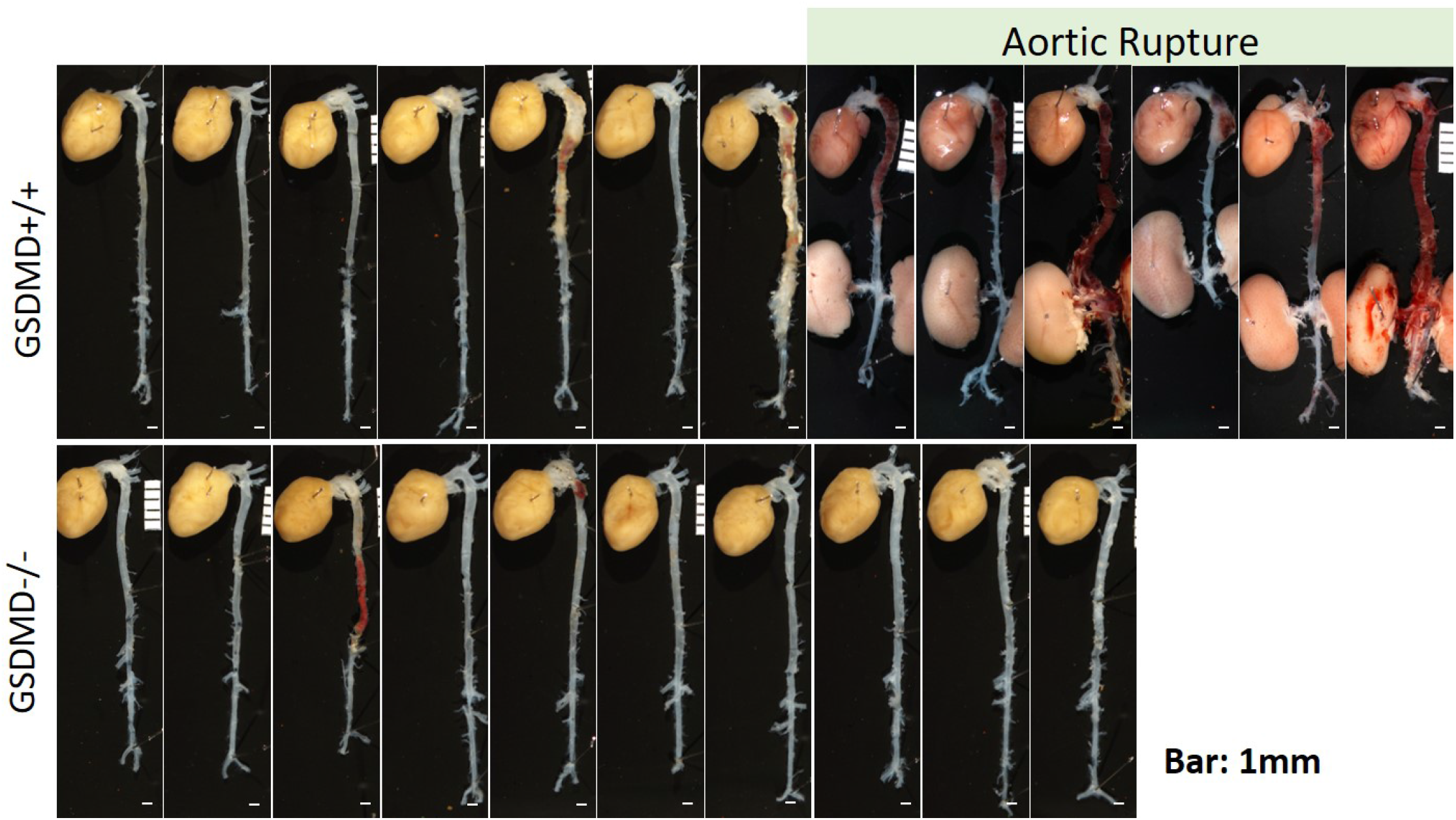
Ex vivo images of aortas from *Gsdmd*^*+/+*^ and *Gsdmd*^*-/-*^ mice administered BAPN.

## Major Resources Tables

### Mouse Models (in vivo studies)

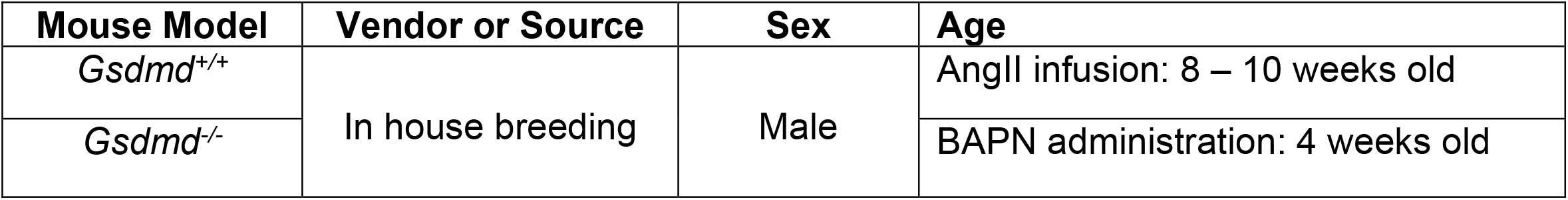

### Mouse Housing Conditions

|  | Mouse Housing Conditions | Note |
| --- | --- | --- |
| Set Temperature range | 22 °C |  |
| Set Humidity range | 50% |  |
| Light : Dark Cycle | 14 hours : 10 hours |  |
| Water | Reverse Osmosis Water | ad libitum |
| Standard Feed | Teklad Irradiated Global 18% Protein Rodent Diet (Envigo); Diet # 2918 | ad libitum |
| Standard Bedding | P.J. Murphy Coarse SaniChip |  |
| Experimental Feed | Envigo; Diet #TD.88137 (AngII study) or Diet # 2918 (BAPN study) | ad libitum |
| SPF | Yes |  |

### Major Reagents/Kits

| Reagents | Vendor or Source | Catalog # |
| --- | --- | --- |
| Angiotensin II | Bachem | 4006473 |
| Beta-aminopropionitrile (BAPN) | MilliporeSigma | A3134 |
| Cholesterol E for plasma total cholesterol concentrations | Fujifilm Wako Diagnostics U.S.A. | 999-02601 |

